# Task success in trained spiking neuronal network models coincides with emergence of cross-stimulus-modulated inhibition

**DOI:** 10.1101/2023.08.29.555334

**Authors:** Yuqing Zhu, Chadbourne M.B. Smith, Mufeng Tang, Franz Scherr, Jason N. MacLean

## Abstract

The neocortex is composed of spiking neuronal units interconnected in a sparse, recurrent network. Neuronal networks exhibit spiking activity that transforms sensory inputs into appropriate behavioral outputs. In this study, we train biologically realistic spiking neural network (SNN) models to identify the architectural changes which enable task-appropriate computations. Specifically, we employ a binary state change detection task, where each state is defined by motion entropy. This task mirrors behavioral paradigms that mice perform in the lab. SNNs are composed of excitatory and inhibitory units randomly interconnected with connection likelihoods and strengths matched to observations from mouse neocortex. Following training, we discover that SNNs selectively adjust firing rates depending on state, and that excitatory and inhibitory connectivity between input and recurrent layers change in accordance with this rate modulation. Input channels that exhibit bias to one specific motion entropy input develop stronger connections to recurrent excitatory units during training, while channels that exhibit bias to the other input develop stronger connections to inhibitory units. Furthermore, recurrent inhibitory units which positively modulated firing rates to one input strengthened their connections to recurrent units of the opposite modulation. This specific pattern of cross-modulation inhibition emerged as the optimal solution when imposing Dale’s law throughout training of the SNNs. Removing this constraint led to the absence of the emergence of this architectural solution. This work highlights the critical role of interneurons and the specific architectural patterns of inhibition in shaping dynamics and information processing within neocortical circuits.

## 1 Introduction

Neocortical networks use neuronal spikes to perform computations which are ultimately responsible for purposeful animal behavior. While definitions of “computation” vary, ranging from precise information calculation to metaphor, one way to study it is as the input / output transformation which underlies the generation of behaviors appropriate to sensory inputs. Through the use of task optimized spiking neural networks (SNN) that were modeled after neocortex, we were able to directly investigate computational mechanisms by comparing the structure and dynamics of models that were or were not executing computations.

SNN models have enriched our understanding of synaptic and cellular mechanisms that underlie neocortical activity in the absence of explicit computations. In particular, they have yielded compelling theories as to the relationships between neocortical structure, dynamics, and computation through measures such as information capacity (e.g. in Brunel 2016), dynamic range (e.g. in Shew et al. 2009), information transmission (e.g. in Mejias & Longtin 2012), and decodability (e.g. in Cohen et al. 2020). For example, asynchronous spiking activity, which is observed in neocortex, has been shown to enhance coding capacity (Kohn et al 2016; Lankarany & Prescott 2015). Recent advances in machine learning using spiking units (Lee et al. 2016; Huh & Sejnowski 2018; Bellec et al. 2020; Zenke & Vogels 2021) enable neuroscientists to apply the same mechanism-uncovering approaches to task-optimized SNNs.

Trained SNNs enable extensive fidelity between model and biology, as they use spike-based computations and learning rules (Brette 2015; Verzi et al. 2018) and can capture biological network nonlinearities (Brette & Gerstner 2005). Spikes are an activity hallmark of neurons within nervous systems, and are crucial in neocortical information processing. The relative timing of spikes has been demonstrated to carry information about stimulus features in neocortex (deCharms & Merzenich 1996). The discrete and sparse nature of spikes can help mitigate the impact of noisy inputs or other fluctuations (Calaim et al. 2022), especially when neuronal units themselves are also leaky (Sharmin et al. 2020). This makes SNNs especially suitable for modeling neural systems operating under realistic, noisy conditions. Individual spikes also have genuine efficacy, as they are used by animals to drive their own behaviors. The earliest stimulus-evoked spikes in mouse V1 are preferentially weighted for guiding behavior (Day-Cooney et al. 2022), and mice are capable of performing a visual discrimination task within a narrow time window during which the majority of V1 neurons involved in the task fire either one or no spikes (Resulaj et al. 2018).

Guided by experimental observations, we built SNNs with realistic neocortical structural and dynamic features and trained them to perform a task analogous to tasks that mice are capable of. SNNs were composed of distinct excitatory and inhibitory adapting units that obeyed Dale’s law, had sparse recurrent connections according to mouse visual cortical statistics (Billeh et al. 2020), long-tailed weight distributions (Song et al. 2005), and low spike rates (Koulakov et al. 2009; Roxin et al. 2011). SNNs were trained with modified backpropagation through time (BPTT; Bellec et al. 2020) to generate a binary report in response to the motion entropy level (low or high) of drifting dot videos.

Over the course of training, we found that networks solved the task by elevating firing rates in response to one level of motion entropy and suppressing firing rates to the other. Specific patterns of connectivity emerged in support of this solution. In particular, recurrent units that have elevated firing to one motion entropy level developed strong inhibitory connections to units that are more responsive to the opposite motion entropy level. The emergence of this pattern of connectivity depended on Dale’s law being imposed throughout training and supports the theory that intracortical inhibitory feedback defines excitatory selectivity in neocortex by refining thalamic inputs (Carandini & Ringach 1997; Katzner et al. 2011).

## 2 Results

### 2.1 Building and Training Biologically Realistic Spiking Neural Network Models

We constructed each recurrent spiking neural network (SNN) model with adaptive leaky integrate and fire (ALIF) model neurons, or “units”. Units are connected to one another via weighted, directed edges which simulate current-based synapses. Networks contained a total of 300 neuronal units; 240 were excitatory (e) and 60 were inhibitory (i), matching the 4:1 e:i ratio observed in neocortex (Figure 1A).

**Figure 1:**
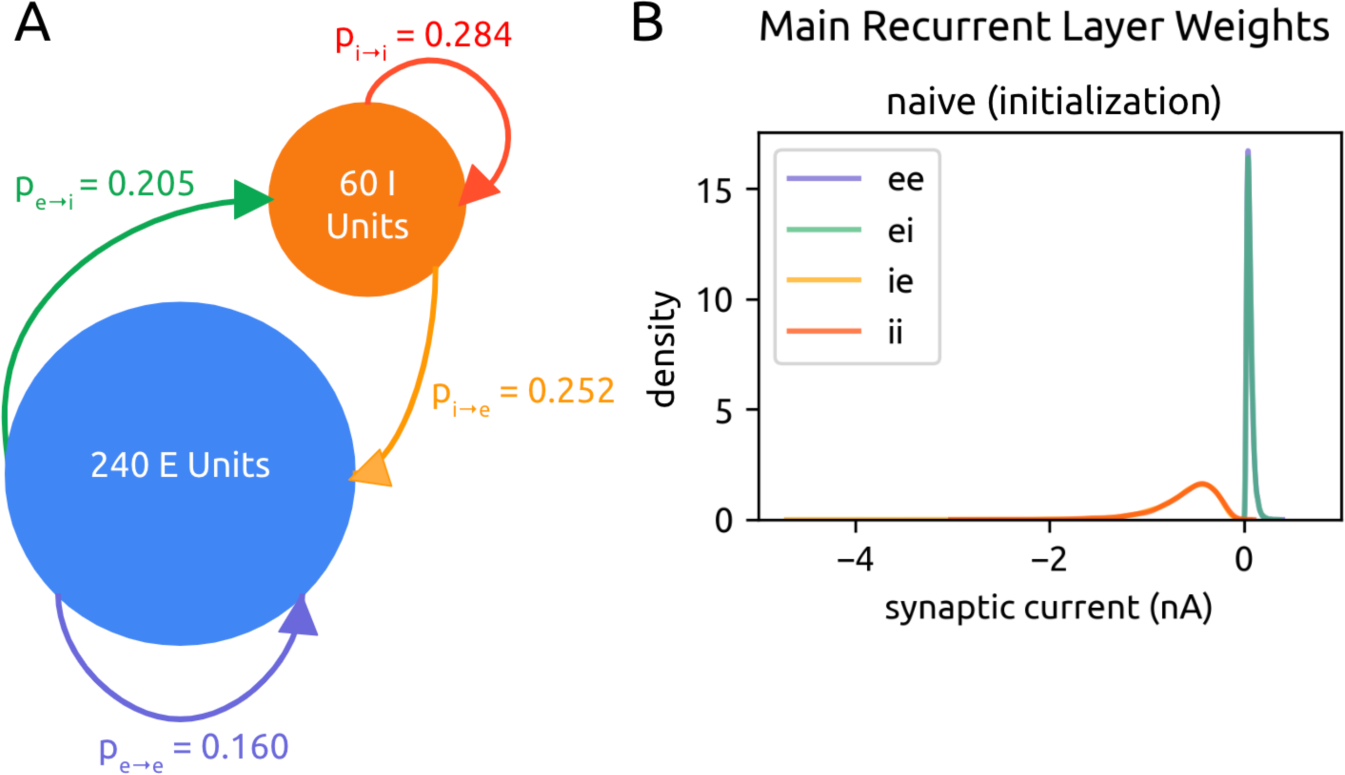
Network architecture. (A) The main recurrent SNN is made of 240 excitatory (E) units and 60 inhibitory (I) units, connected within and between themselves with probabilities of connectivity taken experimentally from mouse neocortex (Billeh et al. 2020). (B) Distribution of naive (upon initialization) recurrent weights of e→e, e→i, i→e, and i→i connections, pooled across all experiments. Inhibitory weights are initialized to be −10x stronger than excitatory weights.

Models were initialized with additional neocortical structural properties. Initial excitatory weights followed a long-tailed distribution (Song et al. 2005), where μ = −0.64 nA, σ = .51 nA, corresponding to a mean of 0.6005 nA and a variance of 0.1071 nA (Bojanek & Zhu et al. 2020) (Figure 1B). Inhibitory weights followed a similar distribution, but with weight values multiplied by −10 (Litwin-Kumar & Doiron 2012). A given excitatory neuron could only have positive outgoing weights, and a given inhibitory neuron could only have negative outgoing weights. Positivity and negativity of edges, i.e. the excitatory or inhibitory identity of each neuron, was maintained throughout training consistent with Dale’s law (Kandel 1957). Each trial had a total duration of 4080 ms. Half of all trials had a change in motion entropy which occurred at a random time between 500 and 3500 ms within the trial. The model is tasked to report the state of motion entropy at each ms time step of the trial instantaneously as output (Figure 2B). The target output sequence is composed of 0’s and/or 1’s (Figure 3). The numerical label assigned to each motion entropy level is randomly swapped in different experiments.

All units were sparsely and recurrently connected; the precise probabilities of connection within and between e and i populations are taken from mouse visual cortex (^p^_e→e_: 0.160, p_e→i_: 0.205, p_i→e_: 0.252, p_i→i_: 0.284, Billeh et al. 2020; Jabri and MacLean 2022) (Figure 1A). Precise connectivity of e and i populations was permitted to change during training. However, overall sparsity was maintained according to the DEEP R algorithm (Bellec et al. 2018), details in Methods.

Models were trained on a visual motion entropy state change detection task, consistent with tasks that mice are able to learn (Douglas et al. 2006; Kirkels et al. 2018; Marques et al. 2018) (Figure 2A). The SNN was presented with spiking output from a model of retina (Maheswaranathan et al. 2023; see below and Methods) evoked by videos of drifting dots moving in one of 4 global directions (0°, 90°, 180°, 270°) and at two levels of motion entropy (high entropy: 15% of dots had the same direction of motion; low entropy: 100% of dots had the same direction of motion).

**Figure 2:**
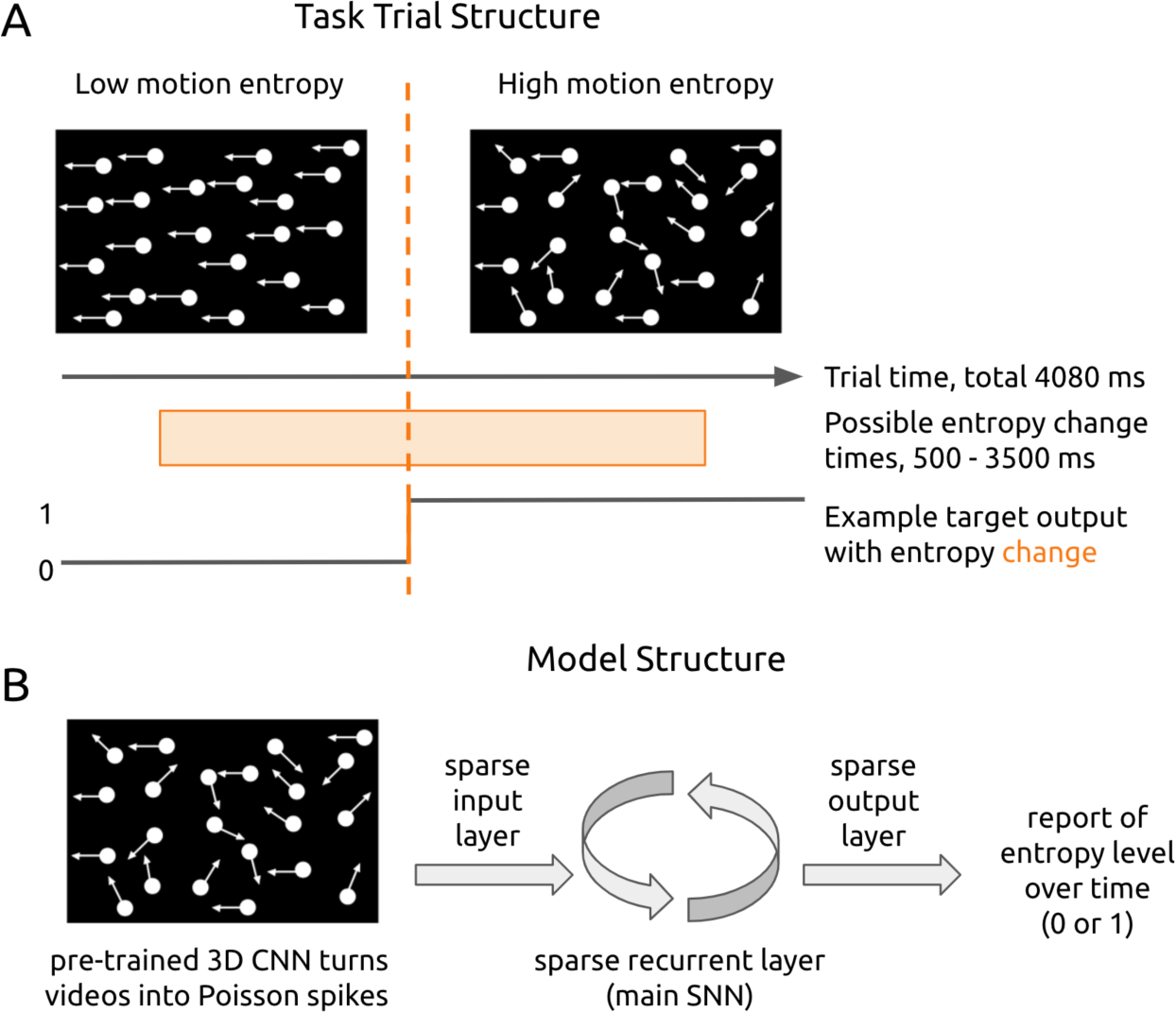
Task and model structure. (A) In each trial, the model is presented with a 4080-ms video of drifting dots. Dots move in 1 of 4 global directions at a high (15% same direction) or low (100% same direction) motion entropy level. The task is to report the dots’ motion entropy level over time. In half of all trials, the entropy level changes at a random time between 500 to 3500 ms. The change can be either from high to low or vice versa. (B) The main SNN receives video input from 16 input channels in the form of Poisson spikes. These 16 input channels convey the activation of a velocity-trained 3D CNN (see Methods and Figure S1) in response to video input. The output of the SNN is a vector of ‘0’s and ‘1’s over time to signal the motion entropy level at each ms. Whether ‘0’ or ‘1’ indicates high entropy or low is toggled in different experiments.

**Figure 3:**
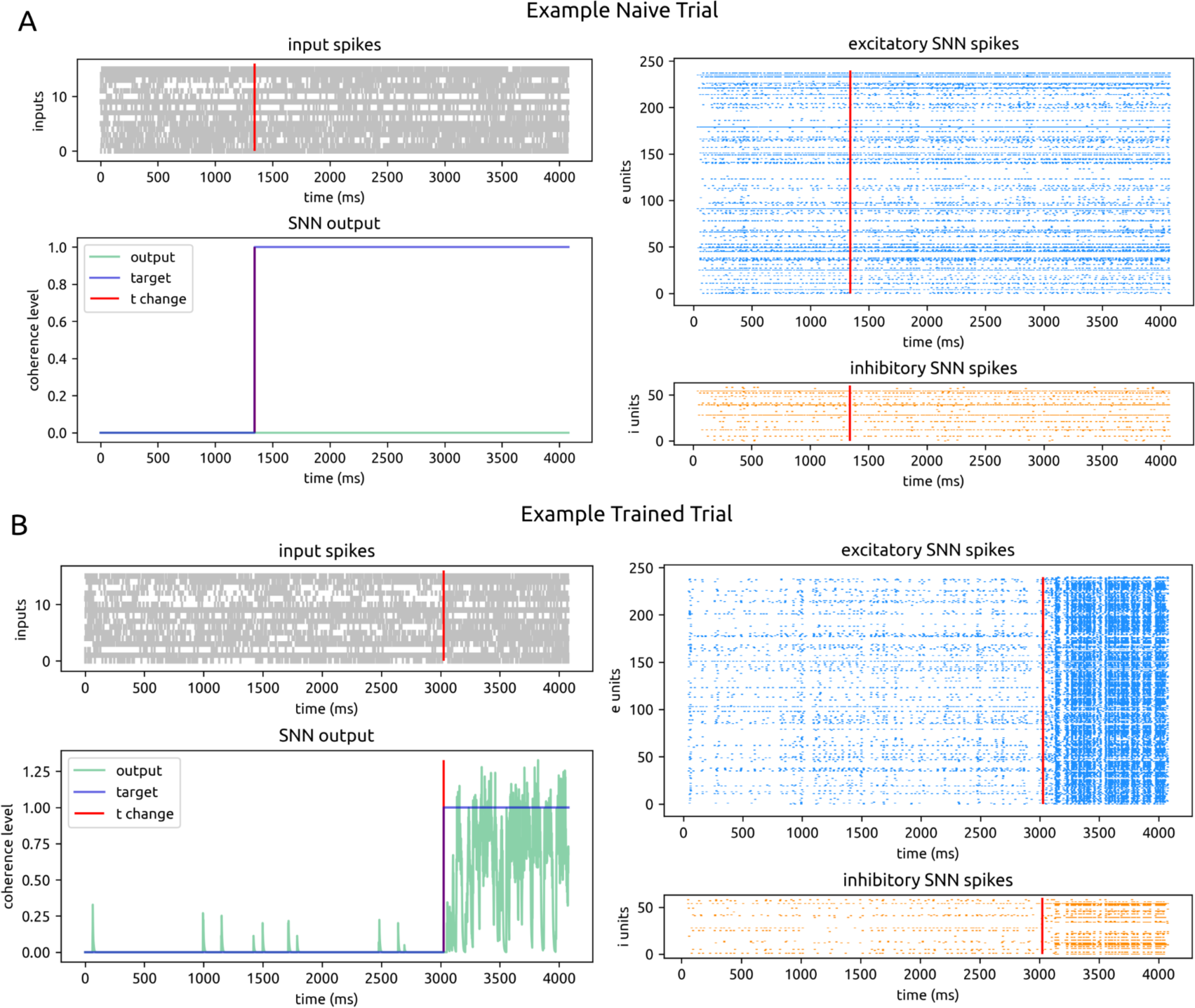
Example model activity. (A) An example of a naive trial in which a motion entropy change occurred. All plots are displayed over time (4080 ms trial duration) on the x-axis. The red vertical line shows the time of motion entropy change. Top left: spikes from 16 input channels. Bottom left: model output for motion entropy level in green; true target motion entropy level in violet. Top right: spikes from 240 excitatory units in the main recurrent SNN. Bottom right: spikes from 60 inhibitory units in the main recurrent SNN. (B) Same as A for an example trained trial. Note the higher firing rates for the motion entropy level ‘1’ and lower rates for ‘0’, which we report on further in section 2.2 and summarize in Figure 6.

To enable the main recurrent SNN to interpret input videos, a 3D convolutional neural network (CNN) model (Maheswaranathan et al. 2023) was used to convert videos into spike sequences (Figure 3, Figure S1A). This CNN was pre-trained to report the velocity (x,y) of global motion in a combined dataset of drifting dot videos and black-and-white natural motion videos (see Methods, Figure S1A). The output activations of the 16 units in the CNN’s next-to-last layer are interpreted as firing rates, from which Poisson spikes were newly generated for each trial. These 16 units were the SNN’s input channels.

The MSE between the model’s motion entropy report and the target motion entropy label (‘0’ or ‘1’) is the task loss. To maintain naturalistic spiking dynamics, a target spike rate was specified (20 Hz, or 0.020 spikes/ms), and the MSE between the recurrent SNN’s actual spike rate from the target rate could be added to the task loss (Zhu et al. 2020). Networks were trained in three ways: to minimize task loss, to minimize rate loss, and to minimize both task and rate loss (Figure 4). We report results primarily for the case of training on both task and rate loss; results from training on only rate and only task loss are reported as controls / points of contrast.

**Figure 4:**
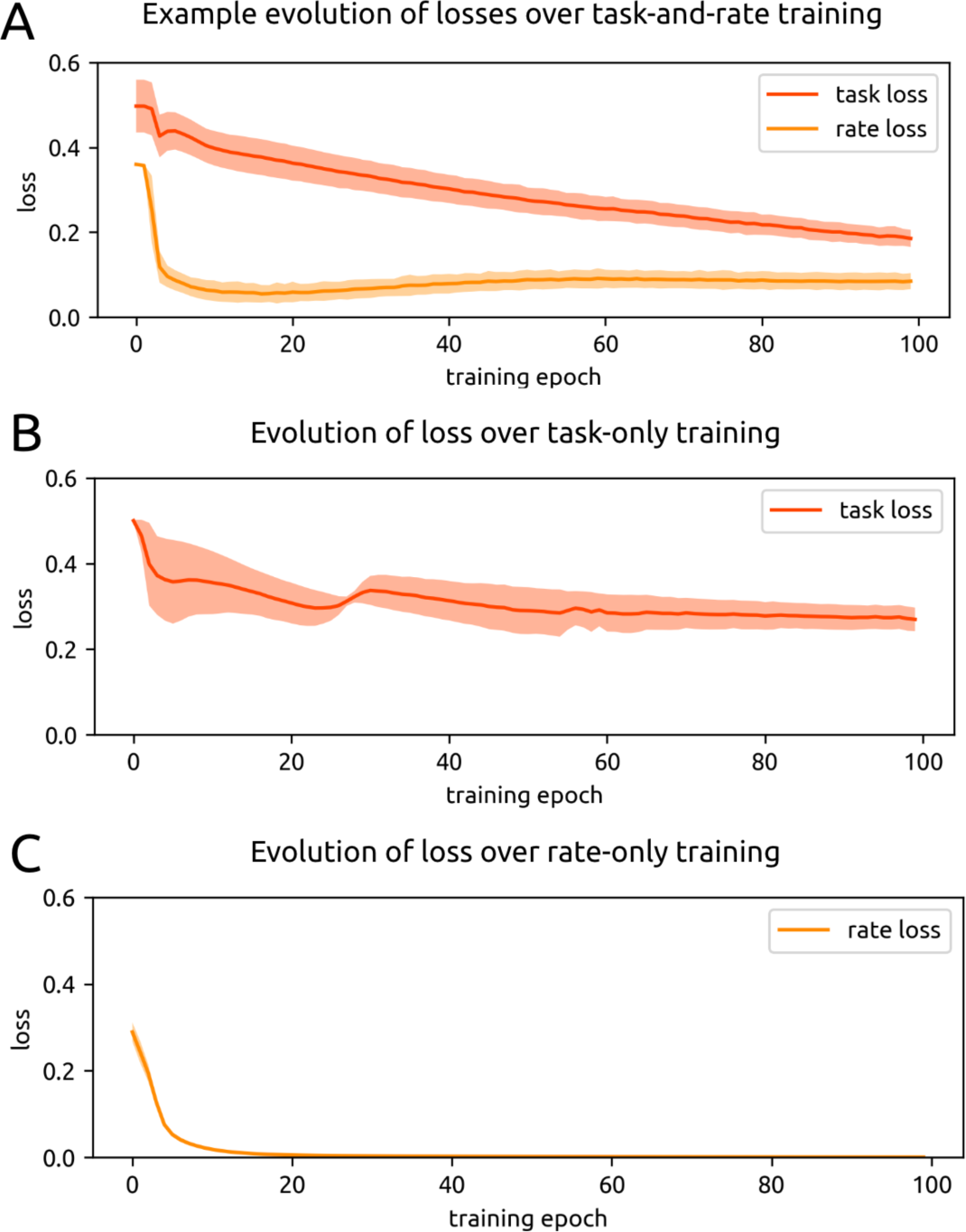
Loss. (A) Losses over training for an example experiment in which the model was trained to minimize both task (red) and rate (orange) loss. Shaded areas are the standard deviation of the losses across trials in each epoch. (B) Loss over training for all experiments in which the model was trained to minimize only task loss. Shaded area is the standard deviation of the epoch task loss across separate experiments. (C) Same as (B) for only rate loss training.

Since networks were all initialized with the same protocol regardless of the class of loss optimization, the naive starting losses across all training sessions (n = 69) were similar, with naive task loss at 0.498 ± 0.073 and naive rate loss at 0.394 ± 0.080.

Across all task-and-rate training sessions (n = 20), models achieved 0.271 ± 0.127 task loss and 0.189 ± 0.174 rate loss after training over 10,000 batch updates, or 100 epochs (Figure 4A). After training on task alone (n = 16), models achieved 0.273 ± 0.021 task loss. After training on rate alone (n = 33), models achieved 0.0007 ± 0.0002 rate loss. Thus training on rate alone leads to better rate performance than when training to minimize both rate and task, but training on task alone does not lead to better task performance than when training to minimize both rate and task loss.

Weights of all layers (input, main recurrent SNN, and output) were permitted to change during each class of training. We used the Adam optimizer, which adapts the learning rate for every variable over the course of training (details in Kingma & Ba 2014). SNNs were trained using backpropagation-through-time (BPTT) with modifications for spiking (see Methods, Huh & Sejnowski 2018; Bellec et al. 2020; Zenke & Vogels 2021).

SNN input and output layers were sparse and matched to the connectivity of the main recurrent and output layers (Figure 5). We also specified that no recurrent units which received input could directly project to output. By stipulating these architectural and training details, we ensured that weight changes which supported training preferentially took place in the recurrent SNN layer, and thus the recurrent SNN layer would play the most important role in solving the task.

**Figure 5:**
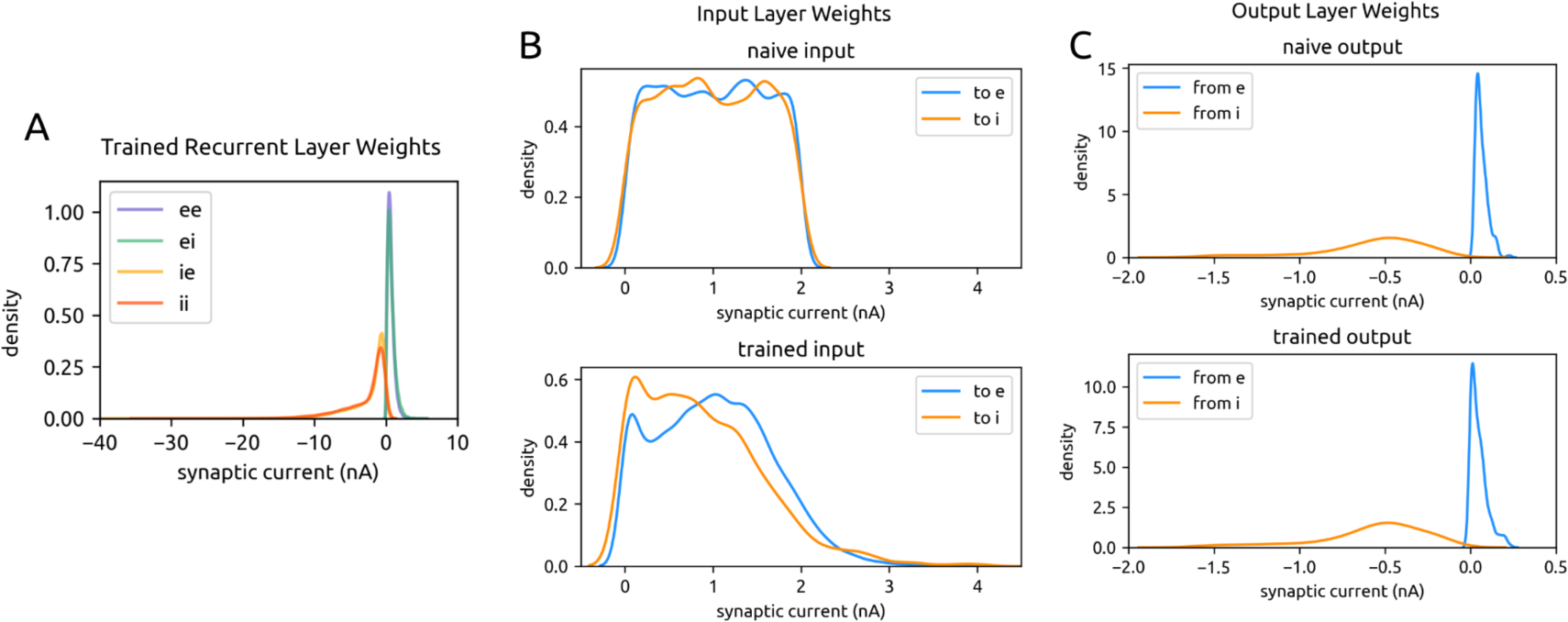
Weight changes over task-and-rate training. (A) Trained weight distributions for the main recurrent SNN. Connections are separated by e→e, e→i, i→e, and i→i type, and weights are pooled across all experiments. Naive recurrent weights are shown in Figure 1B. Note that the long-tailed weight distribution is maintained through training, but scaled roughly 10x. (B) Naive (top) and trained (bottom) input layer weights to main SNN excitatory units (blue) and inhibitory units (orange), pooled across all experiments. Input weights were initialized with a uniform distribution [0, 2] and developed skewness with a longer positive tail over the course of training. (C) Same as A for the output layer; in blue are the weights from main SNN excitatory units and in orange are the weights from inhibitory units. Output layer weights statistically did not change over the course of training.

### 2.2 After Training, Models Demonstrate Significant Synaptic Weight Remodeling in the Recurrent Layer

Synaptic weights are long tailed in neocortex and are initialized as such in our models. This distribution is not enforced during training, yet we observed that a long tailed distribution is preserved following all varieties of training (Figure 1C).

After training on task and rate, weights in the recurrent layer became approximately 10x stronger overall (Figure 5A). Separating the recurrent network by excitatory and inhibitory units, weights change as follows: e→e: 0.010 ± 0.026 naive to 0.113 ± 0.358 trained; e→i: 0.012 ± 0.028 naive to 0.081 ± 0.341 trained; i→e: −0.150 ± 0.307 naive to −0.790 ± 2.131 trained; i→i: −0.169 ± 0.318 naive to −0.768 ± 2.078 trained; p ≈ 0.0 for all naive and trained recurrent weight distributions. Weights in the input layer underwent moderate changes (in→e: 0.325 ± 0.574 naive to 0.432 ± 0.999 trained, p = 2.761 · 10^-152^; in→i 0.288 ± 0.549 to 0.346 ± 0.945 trained, p = 7.923 · 10^-24^) (Figure 5B). Weights in the output layer showed the least difference between naive and trained states; outputs from inhibitory units to outputs do not change at all over training (e→out: 0.010 ± 0.027 naive to 0.006 ± 0.023 trained, p = 6.268 · 10^-15^; i→out: −0.148 ± 0.302 naive to −0.145 ± 0.298 trained, p = 1.0) (Figure 5C). Weight changes for rate-only-training and task-only-training are similar (see Methods). We next investigated whether the edges which began with the strongest naive weights were also the edges with the strongest trained weights. We identified the recurrent edges which had the top decile of initial absolute weights and tracked them over the course of task-and-rate training. After completing training, the percentage of those edges which still contained the top decile of weights was only 3.909 ± 1.377% (compared to 100% if all edges initially in the top decile remained in the top decile). When we broadened to tracking the starting top quartile, after training only 10.028 ± 2.857% of those edges remained in the top quartile. Therefore, the specific initial weights did not pre-determine final weights in the recurrent network.

### 2.3 Models Modulate Firing Rates to Solve the Task

We found that models increasingly modulated their firing rates according to the output label associated with each motion entropy level over the course of training. Network models came to increase firing rates in response to the motion entropy labeled ‘1’ and decreased firing rates to the motion entropy labeled ‘0’ (Figure 3B; Figure 6). It did not matter whether ‘1’ was attached to low or high motion entropy; networks trained with swapped labels exhibited the same behavior, in which the ‘1’ label became associated with elevated firing.

**Figure 6:**
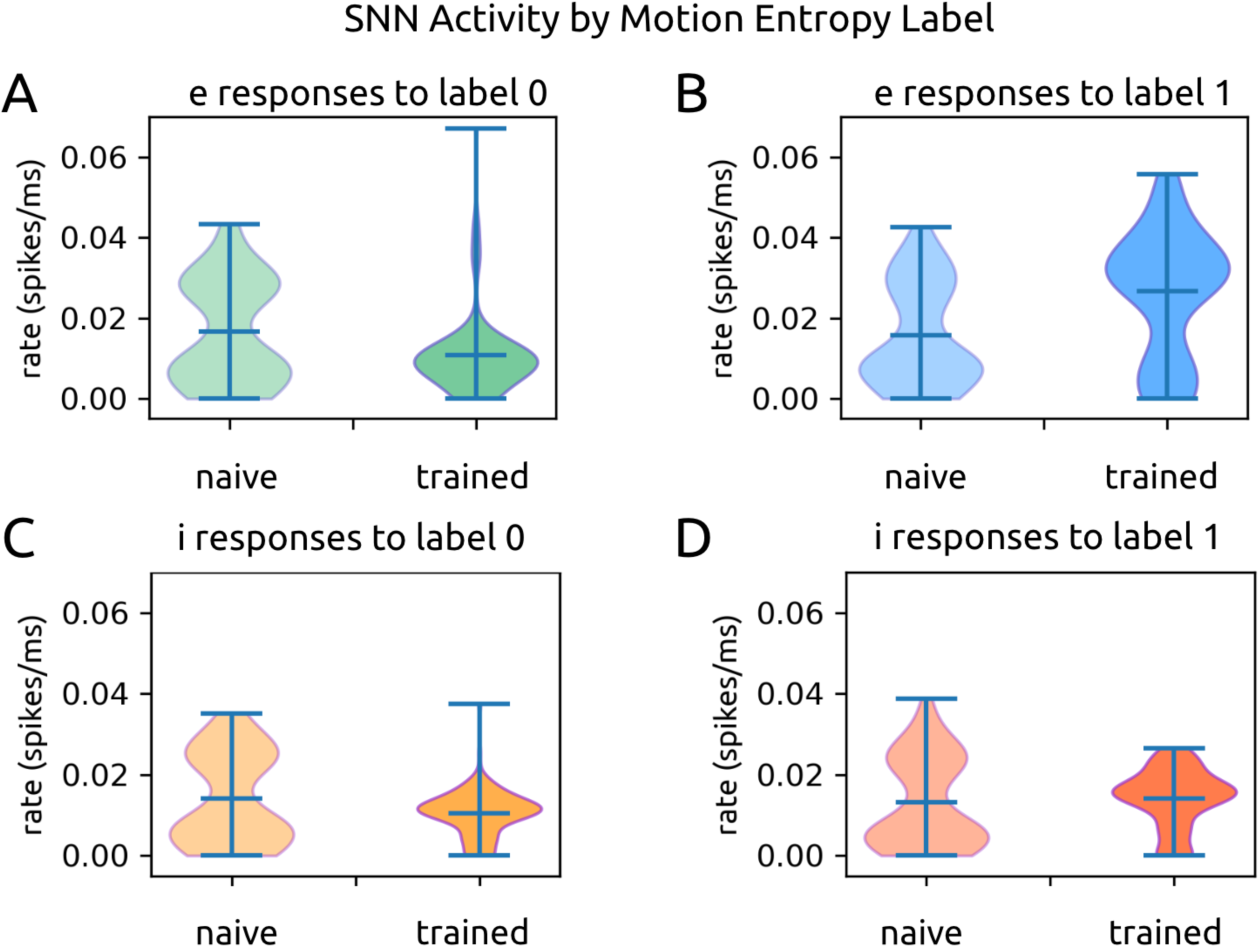
SNN activity in response to motion entropy levels. (A) Excitatory units’ firing rates in response to motion entropy level labeled ‘0’ in the naive (left) and trained (right) states. Rates are pooled over all task-and-rate training sessions. (B) Same as A for entropy level ‘1’. (C) Same as A for inhibitory units. (D) Same as C for entropy level ‘1’.

In the naive state across all training sessions (n = 20), excitatory units responded with 0.017 ± 0.013 spikes/ms to motion entropy ‘0’ and 0.016 ± 0.013 spikes/ms to motion entropy ‘1’, with p = 0.376 comparing responses to the two state labels (Figure 6A;B). We report spiking as spikes/ms since the model reports state on a ms timescale. Inhibitory units responded with 0.014 ± 0.011 spikes/ms to state ‘0’ and 0.013 ± 0.011 spikes/ms to state ‘1’, with p = 0.040 (Figure 6C;D). Thus in the naive state, excitatory and inhibitory units responded similarly to both motion entropy states.

In the trained state (following 10,000 batch updates, or 100 epochs) across all sessions, excitatory units responded with 0.011 ± 0.010 spikes/ms to state ‘0’ and 0.027 ± 0.016 spikes/ms to state ‘1’, p = 6.661 · 10^-16^ (Figure 6A;B). This demonstrates both a decrease and increase in firing rates following training according to label, resulting in an approximately 2.5x increase in firing rates to the ‘1’ state label. Similarly, inhibitory units decreased firing rates to 0.011 ± 0.006 spikes/ms for state ‘0’ and increased firing 0.014 ± 0.007 spikes/ms for state ‘1’, p = 1.147 · 10^-13^ following training (Figure 6C;D). We next investigated how the models’ connectivity changed over the course of training in order to support this reliable behavior and find that both the inputs and the recurrent layer of the SNN change inhibitory connectivity to achieve task accuracy.

### 2.4 Input Channels Strengthen Connections to Excitatory or Inhibitory Recurrent Units According to Their Own Modulation to Motion Entropy Levels

All 16 input channels had different mean firing rates to one state versus the other (all p ≈ 0.0). However, the distance (absolute difference) between channels’ mean responses to one state vs the other was small, measuring 0.021 spikes/ms on average. The difference (high entropy minus low entropy) between mean rates (Figure 7) are as follows for all 16 channels: 0.0163, 0.0002, 0.0192, 0.0874, −0.0176, −0.0163, −0.0063, −0.0170, 0.0284, 0.0033, 0.0180, −0.0395, −0.0336, −0.0118, −0.0166, 0.0071 spikes/ms.

**Figure 7:**
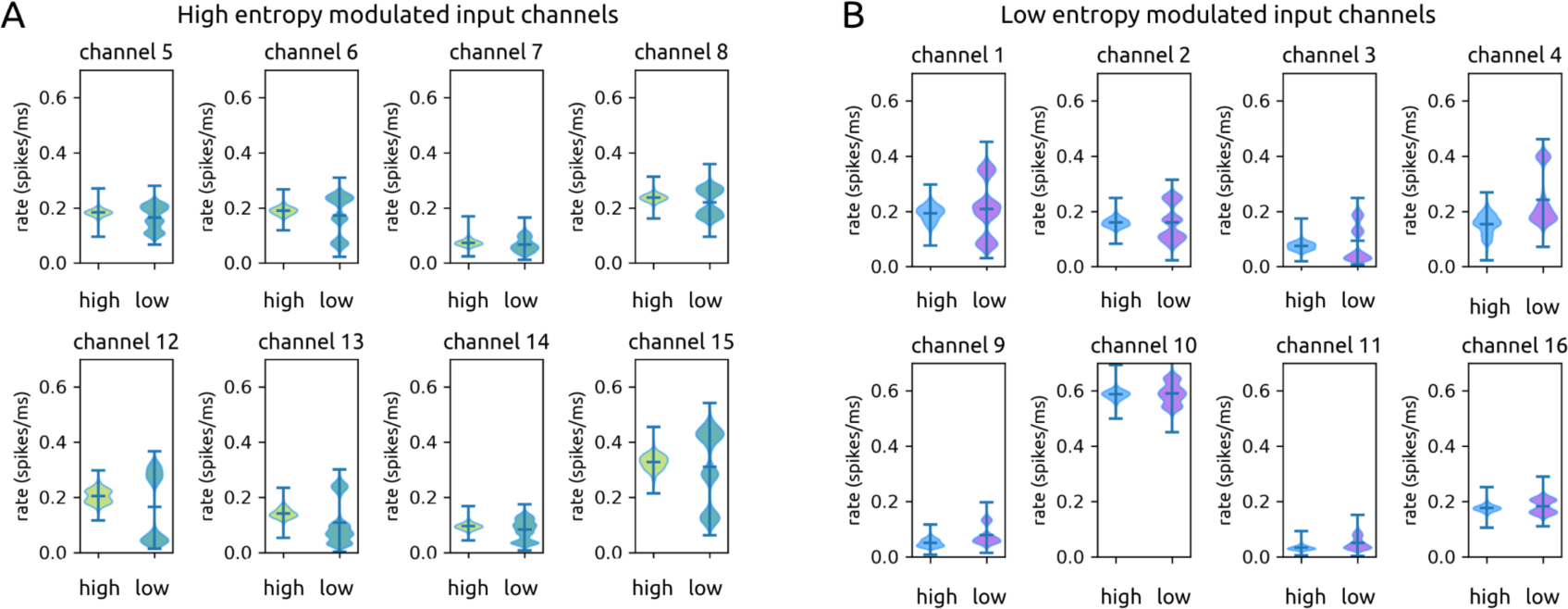
Motion entropy modulation of input channels. (A) Violin plots of input channels’ firing rates in response to high (left violins) and low (right violins) motion entropy levels. A shows the 8 channels that have higher mean firing rates for high entropy. (B) Same as A for the 8 channels that have higher mean firing rates for low entropy.

Notably, half of all channels responded with elevated mean firing during periods of high motion entropy (mean rate to high motion entropy: 0.181 ± 0.078 spikes/ms; rate to low motion entropy 0.162 ± 0.108 spikes/ms; p ≈ 0.0) (Figure 7A). The other half responded with elevated rates to low motion entropy (mean rate to high motion entropy: 0.178 ± 0.165 spikes/ms; rate to low motion entropy: 0.201 ± 0.160 spikes/ms; p ≈ 0.0) (Figure 7B) regardless of label. We will refer to these input channels as high-motion-entropy-modulated and low-motion-entropy-modulated input channels respectively.

Input channels were either positively modulated by high motion or low motion entropy input, but in the case of low motion entropy, input channels exhibited multimodal rate distributions. In contrast, in response to high motion entropy, the input channels’ rate distributions were largely unimodal. When we consider medians instead of means, 10 input channels had greater median rates in response to high motion entropy, and 6 input channels had greater median rates in response to low motion entropy. The average distance between all medians was 0.023 ± 0.029 spikes/ms. When we consider modes, 5 input channels had greater modes in response to high motion entropy, and 6 input channels had greater modes in response to low motion entropy. Finally, 5 input channels had the same modes to both low and high motion entropy, further illustrating the similarity of input responses to the two motion entropy states. The average distance between all modes was 0.045 ± 0.048 spikes/ms. We chose to use the mean as the measure of central tendency to define whether input channels were positively modulated by high or low motion entropy; the majority of low motion entropy rate distributions were trimodal, making the mean the best measure for comparison across all distribution shapes.

Each input channel synapses onto subpopulations of units in the recurrent layer, including both excitatory and inhibitory units. In the naive state, high motion entropy modulated input channels and low motion entropy modulated input channels had similarly weighted connections onto both excitatory (high entropy: 0.198 ± 0.009; low entropy: 0.204 ± 0.004) and inhibitory units (high entropy: 0.187 ± 0.010; low entropy: 0.172 ± 0.011) in the recurrent layer. Over the course of training, the two classes of motion entropy modulated input channels diverged in the strength of their synapses onto excitatory and inhibitory recurrent units (Figure 8A). The recurrent layer, where the majority of training related changes occurred, then amplifies this difference, as we report in a later section.

**Figure 8:**
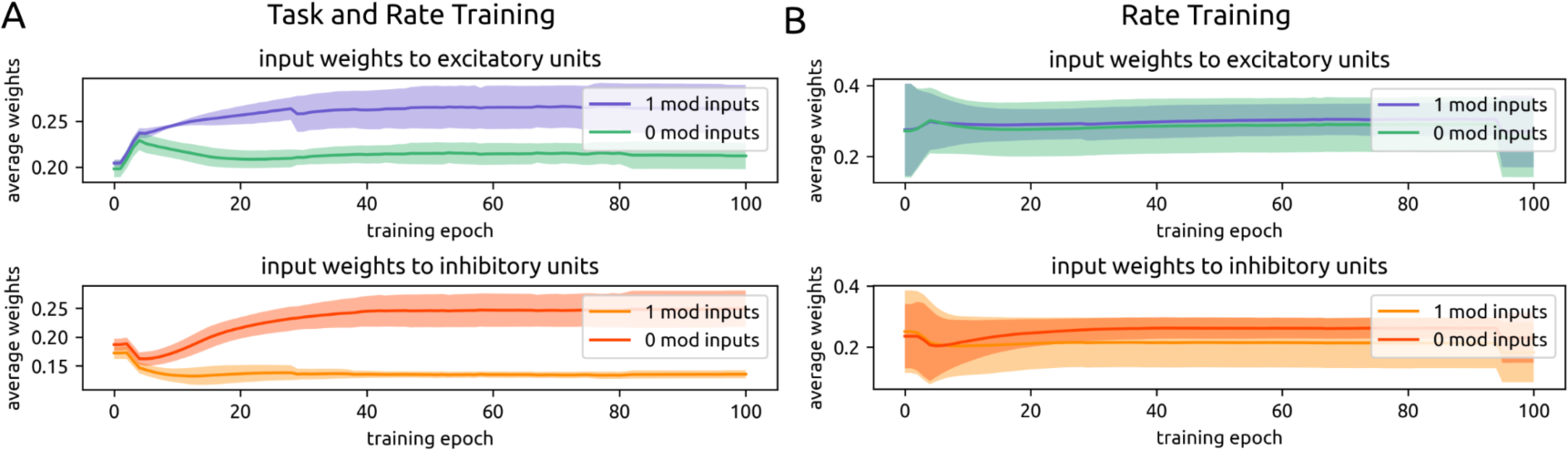
Selective strengthening of input layer weights according to input modulation. (A) Over the course of task-and-rate training, input channels positively modulated by motion entropy 1 develop stronger connections to excitatory units (top), while input channels positively modulated by motion entropy 0 develop stronger connections to inhibitory units (bottom). (B) This pattern is not observed during only rate training.

When high motion entropy is labeled as output ‘0’ and low motion entropy is labeled as output ‘1’ during training, high entropy modulated input channels increase weights onto inhibitory units (0.249 ± 0.032) while decreasing weights onto excitatory units (0.212 ± 0.014). On the other hand, low motion entropy modulated input channels develop stronger weights onto excitatory units (0.263 ± 0.027) and weaker weights to inhibitory units (0.135 ± 0.007). In this way, when a high entropy input is presented, the network becomes inhibition dominated, leading to lower spike rates and a report of the lower output label (‘0’). Conversely, when a low motion entropy input is presented, the network becomes driven by excitation and reports the higher output value (‘1’).

If the labels are swapped, the opposite changes in connectivity occurred. High motion entropy modulated input channels developed stronger weights onto excitatory units and low motion entropy modulated input channels developed stronger weights onto inhibitory units. Thus, over the course of training, input channels that preferentially responded to a particular motion entropy level–regardless of the magnitude of difference– became more strongly connected to either excitatory or inhibitory units, resulting in either high or low firing states to match the desired output label.

This result can be summarized as a ratio of 1-modulated input weights to 0-modulated input weights onto inhibitory and excitatory SNN units. Again, 0 can refer to either low or high motion entropy; what is important is which particular motion entropy is given the ‘0’ label for that experiment. For inhibitory units, this ratio begins at 0.921 in the naive state and becomes 0.543 after training. This means that through training, inhibitory units receive approximately twice as much drive from input channels modulated by the entropy state labeled as ‘0’ in a given experiment. For excitatory units, this ratio begins at 1.031 and becomes 1.239 after training. Thus following training, excitatory units receive moderately more input drive from 1-modulated input channels.

In versions of the model trained only on rate, we do not observe this separation of input channels onto excitatory or inhibitory recurrent populations (Figure 8B). For inhibitory units, the average ratio of 1-modulated input weights over 0-modulated input weights begins at 0.970 and becomes 0.821. For excitatory units this ratio begins at 1.010 and becomes 1.054. This is expected, as there is no imperative to drive inhibition or excitation more strongly in order to report ‘1’ or ‘0’ when optimizing for rate loss alone.

The input layer is not the only part of the model that changes to enable rate modulation–the recurrent layer itself plays a role in amplifying this behavior.

### 2.5 Recurrent Inhibitory Units Strengthen Connections to Recurrent Units of Opposite Modulation Patterns

Similar to input channels, recurrent units also modulated firing rates to one or the other motion entropy level. We defined the positive modulation of recurrent units as the motion entropy label to which units responded with greater mean firing rates in the network’s final trained state.

In the task-and-rate-trained SNN, the majority of excitatory units respond with elevated mean firing to the motion entropy level labeled as ‘1’ (193.8 ± 60.0 out of 240 total excitatory units). A smaller proportion (27.1 ± 15.7 excitatory units) respond with elevated firing to the motion entropy labeled as ‘0’. Similar numbers of inhibitory units respond with elevated firing to ‘1’-labeled (21.6 ± 7.5 out of 60 total inhibitory units) as to ‘0’-labeled (21.3 ± 8.2 inhibitory units) motion entropy input.

We plotted the mean weights within and between ‘0’ and ‘1’ label modulated populations for e→e, e→i, i→e, and i→i connections over the course of task-and-rate training (Figure 9).

**Figure 9:**
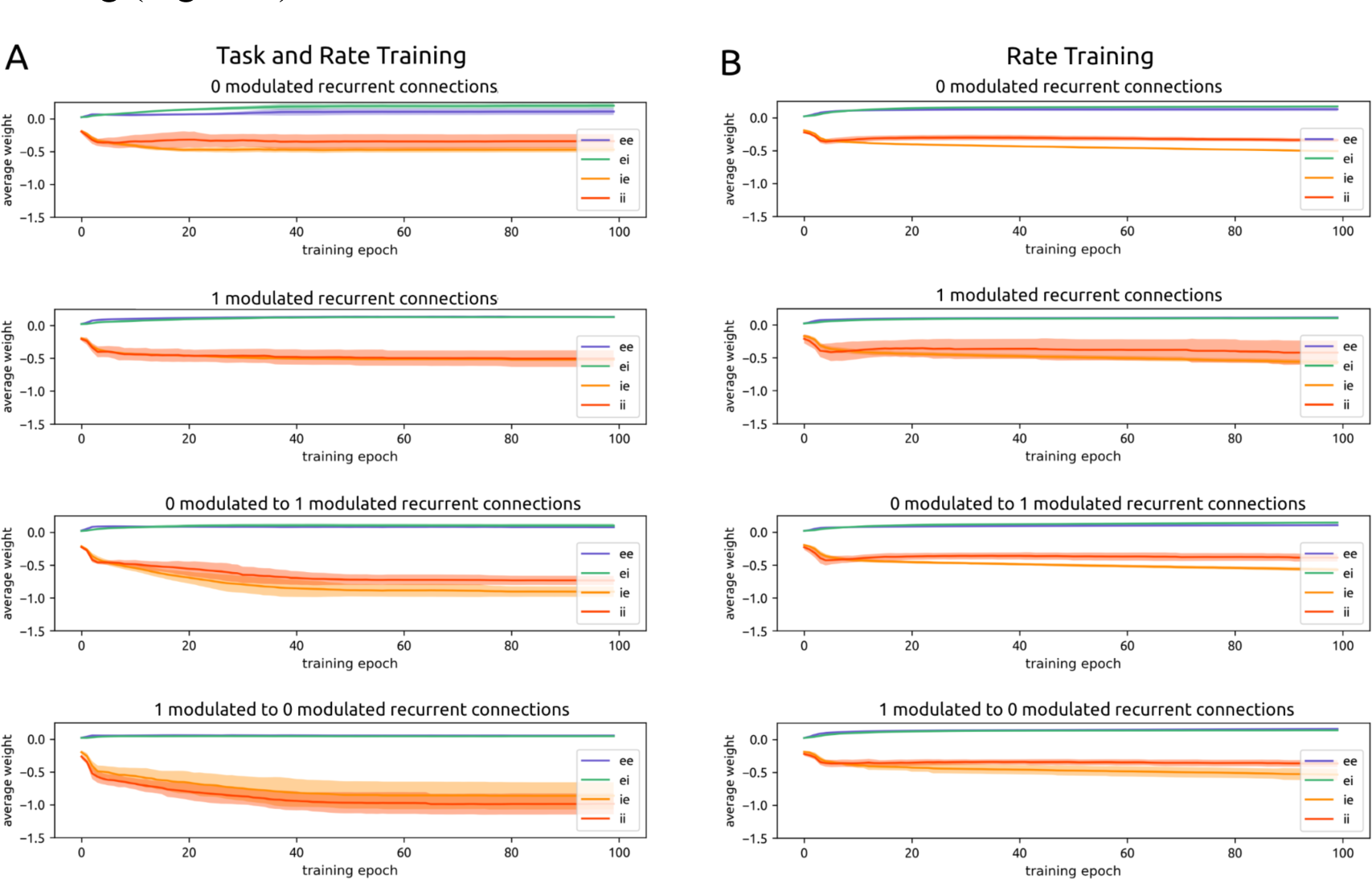
Selective strengthening of recurrent cross-modulation inhibition. (A) Over the course of task-and-rate training, inhibitory recurrent units develop the strongest connections to recurrent units of the opposite modulation pattern (bottom two plots). (B) This pattern is not observed when models are trained on only rate.

Again, these populations are defined according to the label for which they positively modulated firing rates at the end of training. Plots track retrospectively how these units’ weights evolved over training in conjunction with the emergence of positive rate modulation. We found that differences arise in the connectivity between recurrent units according to the units’ final label preference. In particular, recurrent inhibitory units send the strongest connections to units of the opposite modulation to themselves (Figure 9A).

Connectivity for excitatory units to other excitatory units is as follows: ‘0’-modulated to ‘0’-modulated: 0.021 ± 0.007 naive to 0.104 ± 0.052 trained; ‘1’-modulated to ‘1’-modulated: 0.025 ± 0.009 naive to 0.128 ± 0.011 trained; ‘0’-modulated to ‘1’-modulated: 0.024 ± 0.007 naive to 0.076 ± 0.015 trained; ‘1’-modulated to ‘0’-modulated: 0.022 ± 0.007 naive to 0.056 ± 0.013 trained. Over training, recurrent excitatory weights increase more within the same modulation pattern (approximately 5x relative to naive) than across (approximately 3x relative to naive).

Connectivity for excitatory units to other inhibitory units is as follows: ‘0’-modulated to ‘0’-modulated: 0.030 ± 0.014 naive to 0.191 ± 0.046 trained; ‘1’-modulated to ‘1’-modulated: 0.027 ± 0.012 naive to 0.125 ± 0.020 trained; ‘0’-modulated to ‘1’-modulated: 0.025 ± 0.010 naive to 0.097 ± 0.033 trained; ‘1’-modulated to ‘0’-modulated: 0.024 ± 0.010 naive to 0.045 ± 0.004 trained. Once again, over training, excitatory weights onto inhibitory units increase more within the same modulation pattern (approximately 5x) than across (approximately 3x).

Connectivity for inhibitory units to excitatory units is as follows: ‘0’-modulated to ‘0’-modulated: −0.226 ± 0.075 naive to −0.523 ± 0.102 trained; ‘1’-modulated to ‘1’-modulated: −0.243 ± 0.090 naive to −0.605 ± 0.138 trained; ‘0’-modulated to ‘1’-modulated: −0.266 ± 0.100 naive to −0.939 ± 0.158 trained; ‘1’-modulated to ‘0’-modulated: −0.267 ± 0.118 naive to −0.978 ± 0.224 trained. In contrast to excitatory units, over training, inhibitory weights onto excitatory units increase moderately within the same modulation pattern (approximately 2x) and increase more strongly across modulation (approximately 4x).

Connectivity for inhibitory units to other inhibitory units is as follows: ‘0’-modulated to ‘0’-modulated: −0.233 ± 0.067 naive to −0.411 ± 0.149 trained; ‘1’-modulated to ‘1’-modulated: −0.235 ± 0.059 naive to −0.519 ± 0.138 trained; ‘0’-modulated to ‘1’-modulated: −0.280 ± 0.106 naive to −0.855 ± 0.248 trained; ‘1’-modulated to ‘0’-modulated: −0.327 ± 0.135 naive to −1.097 ± 0.266 trained. Over training, inhibitory units moderately increase weights to other inhibitory units of the same modulation pattern (approximately 2x) and more strongly increase weights to other inhibitory units of the opposite modulation pattern (approximately 3x).

Thus, over the course of training, excitatory units increase weights to other units of the same modulation pattern, while inhibitory units increase weights to units of the opposite pattern. Two-sample KS-testing confirms that the trained weight distributions are different for within- and across-modulation excitatory connections (p = 3.205 · 10^-7^ comparing e→e within- vs. across-modulation; p = 3.847 · 10^-10^ comparing e→i within-vs. across-modulation). This is also true for within- and across-modulation inhibitory connections (p = 6.323 · 10^-14^ comparing i→i within-vs. across-modulation; p = 5.409 · 10^-11^ comparing i→e within-vs. across-modulation).

This set of results can be summarized as a ratio: mean weight across-modulation / mean weight within-modulation. For inhibitory connections in naive networks, this ratio is 1.215. After training, this ratio increases to 1.879. Thus inhibitory units in trained networks send connections that are approximately twice as strong to opposite-modulated recurrent units compared to same-modulated recurrent units. For excitatory connections, this ratio is 0.928 in the naive state and 0.500 in the trained state. Thus excitatory units exhibit the precise opposite pattern of connectivity as inhibitory ones after training: they come to send connections that are twice as strong to *same-*modulated recurrent units compared to opposite-modulated recurrent units.

When networks are trained on rate alone (n = 38), this pattern of stronger cross-modulation inhibition is not observed (Figure 9B). The average ratio of inhibitory weights across-modulation / within-modulation is 1.035 for naive networks and 1.027 for trained networks. For excitatory weights, the ratio is 0.996 for naive networks and 1.051 for trained networks. The proximity of this ratio to 1 informs us that the strength of across-modulation vs. within-modulation connections are approximately equal in strength for the rate-trained network.

### 2.6 While the Specifics of Inhibition are Not Crucial for the Development of Cross Modulation Inhibition, Adherence to Dale’s Law is Required

Through their strong cross-modulation connectivity, inhibitory units enable the appropriate modulation of high and low recurrent firing rates in response to ‘1’ and ‘0’ labeled motion entropy states. However, inhibitory weights are initialized to be 10x stronger than excitatory weights; it is possible that this initialization predisposes the network to reach this solution. We therefore initialized networks (n=21) with recurrent inhibitory weights that were only slightly (1.5x) stronger than excitatory weights (Figure 10C).

**Figure 10:**
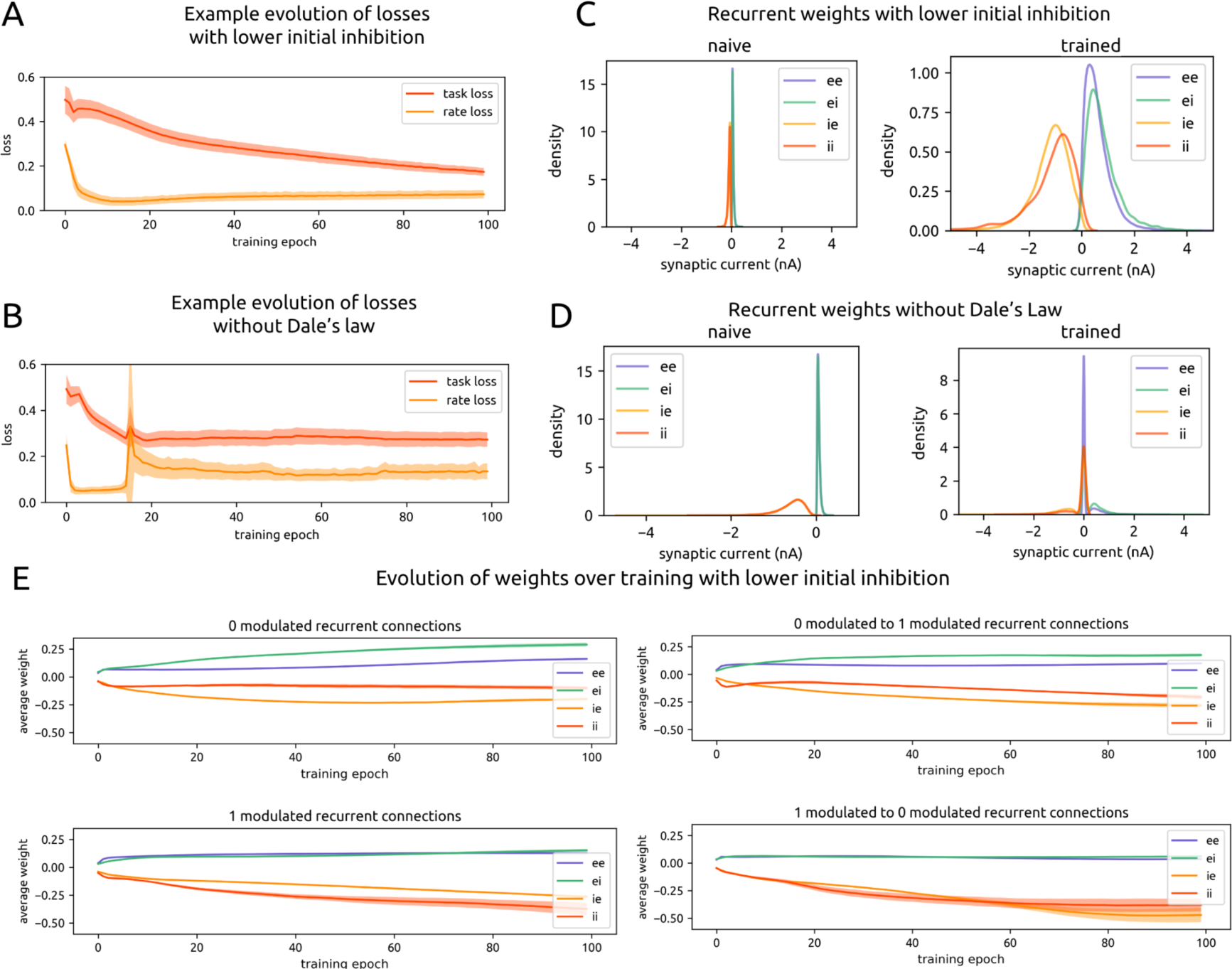
Relaxing constraints on inhibition. (A) Task and rate loss over the course of training for an example experiment in which the model was initialized with lower inhibitory weights (1.5x as strong as excitatory weights, compared to 10x previously). (B) Same as A for a model without Dale’s law constraints. (C) Naive (left) and trained (right) weight distributions for all models with lower initial inhibition. Note that while all weights do increase, the 1.5x relationship is largely preserved. (D) Same as C for all models trained without Dale’s law constraints. Note that the designation of units as initially excitatory or initially inhibitory does not mean anything for the negativity or positivity of weights after training. (E) Same as Figure 9A for all models with lower initial inhibition.

Following successful training on task and rate loss (Figure 10A) in the same manner as before, we found that rate modulation according to motion entropy label persisted (‘1’ mean firing rate: 0.030 ± 0.004 spikes/ms for e units and 0.021 ± 0.005 spikes/ms for i units; ‘0’ mean firing rate: 0.013 ± 0.002 spikes/ms for e units and 0.016 ± 0.002 spikes/ms for i units).

Overall trained weight distributions (Figure 10C) differed from when inhibitory units were initialized to be 10x stronger. Excitatory weights were stronger than before (trained e→e: 0.643 ± 0.633 now, previously 0.113 ± 0.358; trained e→i: 0.802 ± 0.609 now, previously 0.081 ± 0.341). However, trained inhibitory weights did not increase as much; they approximately preserved the 1.5x magnitude difference from excitatory weights with which they were initialized (trained i→e: −1.193 ± 0.700 now, previously −0.790 ± 2.131; trained i→i: −1.148 ± 0.904 now, previously −0.768 ± 2.078).

When we examined the evolution of recurrent connectivity over the course of training, we found that inhibitory cross-modulation persisted (Figure 10E). The ratio of mean inhibitory weights between oppositely-modulated units over similarly-modulated units was 1.121 in the naive state and 2.002 after training. This reflects the two-fold strengthening of inhibitory weights across modulation type versus within modulation type, as seen previously when inhibitory weights were initialized to be stronger. For excitatory weights, this ratio was 0.918 in the naive state and 0.466 in the trained state, reflecting that excitatory weights once again become stronger between units of the same modulation type. Thus, despite changes in the overall weight distributions upon initialization and in the trained state, the pattern of stronger cross-modulation inhibition and stronger within-modulation excitation still arose as a task solution.

We next investigated the extent to which this pattern of cross-modulation inhibition is preserved in models (n=19) when, in addition to weakening of initial inhibitory weights, Dale’s Law was removed as a constraint during training. In particular, initial excitatory units could develop negative outgoing weights, and initial inhibitory units could develop positive outgoing weights. Under these further relaxed conditions, would cross-modulation inhibition still emerge as a preferred task solution?

Following training in the same manner as before (Figure 10B), weight distributions were no longer meaningfully separated into excitatory or inhibitory according to the identity of the presynaptic unit (Figure 10D). The magnitudes of all positive and negative weighted edges remained comparable to those of the naive state (excitatory: 0.060 ± 0.033 naive to 0.119 ± 0.281 trained; inhibitory: −0.090 ± 0.049 naive to −0.063 ± 0.264 trained), whereas formerly, under Dale’s law constraints, all weights increased approximately 10x over the course of training (Figure 1B;5A).

We quantified the extent of cross-modulation inhibition with the same ratio as before, which is that of mean weights across-modulation over mean weights within-modulation. Because we could no longer track units and their edges according to excitatory and inhibitory identity, we report this ratio for the trained model only. For all negative edges, this ratio was 0.842 in the trained state. For all positive edges, this ratio was 0.914 in the trained state. The proximity of both ratios to 1 indicates that, unlike in previous versions of the model, the model without Dale’s law did not heavily rely on cross-modulation inhibition and within-modulation excitation to solve the task. We hypothesize that, since each neuron’s activity could now directly drive both downstream excitation and inhibition, the model developed more specific connectivity towards a solution. These specific connections between individual units could no longer be summarized through excitatory and inhibitory population groups. It is also notable that the model did not achieve as robust a task performance level as previously (Figure 10B compared to Figure 4A); this may be due to the challenges of needing to develop hyper-specific connections in order to balance the dual effects of excitation and inhibition from individual neurons.

## 3 Discussion

Using biologically realistic, task-optimized SNN models, we found that recurrent networks of spiking units selectively elevate or depress firing rates in response to a motion entropy-report task. We achieved a level of interpretability, finding that excitatory and inhibitory connectivity of the input and main recurrent layers changed in conjunction with this rate modulation.

Input channels that responded preferentially to the motion entropy level labeled as ‘1’ (numerical labels attached to low or high motion entropy were swapped in certain experiments) developed stronger connections to excitatory units in the main recurrent SNN. In contrast, input channels that preferred motion entropy level ‘0’ strengthened connections to inhibitory units. This was the first step leading to excitation-dominated or inhibition-dominated SNN activity. This effect was further amplified through connectivity patterns in the main recurrent SNN. Recurrent inhibitory units that were positively modulated by one motion entropy label at the end of training demonstrated a strengthening of connections to both excitatory and inhibitory units of the opposite modulation. Thus when one motion entropy level was presented, inhibitory units would suppress other units that were not part of the appropriate motion entropy modulation. This arose over the course of training. In such a way, small but significant differences in the input statistics are exploited, and the recurrent network employs strong cross-modulation inhibition to further adjust its responses to be appropriate to the input.

A biological parallel to the emergence of this pattern–in which units positively modulated by one motion entropy level came to inhibit units of opposite modulation–can be found in the rich literature of cross-orientation suppression of V1 neurons (Morrone et al. 1982; Eysel et al. 1990; DeAngelis et al. 1992; among others) and of the role of interneurons in establishing V1 excitatory selectivity to stimulus orientation and direction. While the relative contributions of thalamic (feedforward) and intracortical (feedback) connections to establishing V1 selectivity is under continued study (Ferster & Miller 2000; Alitto & Dan 2010; Katzner et al. 2011), one theory is that selectivity arises through a combined effect of initially broad selectivity via LGN inputs that is then sharpened through intracortical feedback (Carandini & Ringach 1997). Our modeling results support this theory in that the strict dichotomy between thalamic and intracortical contributions is false. We find that intrinsic modulation patterns from our input channels, which can be interpreted as modeling thalamic inputs, was refined by recurrent connections in our main SNN, which can be interpreted as a model of V1. As task learning progressed, the local recurrent circuitry underwent changes to enhance the amplification of input modulation patterns by strengthening recurrent excitatory connections from like-modulated excitatory units and cross-modulated inhibitory units. This is the ideal architectural solution in the presence of Dale’s law. Removal of Dale’s law as a constraint eliminated this as an optimal solution and also degraded performance of the models.

Intracellular data and modeling work suggests that this local intracortical feedback takes the form of inhibitory connections from interneurons that have the same–rather than broader–selectivity as excitatory targets (Katzner et al. 2011). By matching excitatory selectivity, interneurons can precisely keep responses to undesired stimuli below the spike threshold. This is supported by the variety of connectivity according to modulation observed in our trained models, and the mechanisms underlying this behavior are aligned in both neocortex and the model, since the activity of our model neurons is also fundamentally based on thresholding.

A potential candidate responsible for establishing excitatory selectivity in neocortex is parvalbumin-expressing (PV) interneurons. PV neurons synapse onto the soma or axon of synaptic partners (Kepecs & Fishell 2014), allowing them to tightly control spiking in postsynaptic neurons. We used a generic inhibitory neuron in our models, but this property of direct control is a feature of our inhibitory model units. This property makes PV neurons a candidate for selecting neurons that are involved in task-relevant assemblies. Theoretical work suggests that PV neurons stabilize new groups of task-associated excitatory neurons (Bos et al. 2020, Lagzi et al. 2021), which is consistent with experimental work in associative learning (Morrison et al. 2016).

While there were various possible network configurations that could have achieved stimulus modulation, such as strengthening excitatory connections, it is noteworthy that our networks converged on this specific pattern of cross-modulation inhibition through training. This result may underscore the significance of PV neurons and their connectivity pattern in establishing local circuit computations in the neocortex. A natural extension of this work is to diversify neuronal subtypes in future models, as each exhibits unique connectivity properties in local circuits, which may point to distinct computational roles as well (Kepecs & Fishell 2014; Cone et al. 2019).

Through this work, we provide two circuit mechanisms–one based on input and another based on recurrent connectivity changes–that underlie task learning. Moreover, we achieve this result in a spiking network, making the circuit mechanisms more directly applicable to networks of spiking neurons in neocortex. We believe that task-trained SNNs demonstrate promise for advancing the understanding of network computation and applying that understanding to neocortex, and look forward to future work that adopts this method for this goal.

## 4 Methods

### 4.1 Model construction

The models we are building and training are recurrent spiking neural networks (SNNs). Each model is built with spiking units which represent individual neurons. Units are connected with one another via weighted, directed edges which represent synapses. The weight of an edge 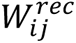 is a numerical value that indicates the strength of the connection from presynaptic unit *i* to postsynaptic unit *j* within the recurrent SNN.

In a network of 300 neuronal units, 240 are excitatory and 60 are inhibitory. They are built this way to maintain the 4:1 e:i ratio, which is a ratio observed in neocortex. All synaptic edges originating from an excitatory unit have positive weight values, and all from an inhibitory unit have negative weight values. Positivity and negativity of each edge is maintained during training, although values may change (Zhu et al. 2020).

### 4.2 Input

Input to the main SNN will vary depending on the task of interest. The task and input preprocessing are described in detail in the “Task” section. As an overview, input is delivered in the form of spike sequences onto a subset of the main SNN units. Input spikes are also weighted (e.g. 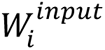 is the input weight to SNN unit *i*), and the weight distribution can be initialized as desired and permitted to change or be fixed during training. For the majority of our experiments, input weights were initialized with a sparse uniform distribution (*min* = 0.0, *max* = 0.4) and all weights were permitted to change during training while overall sparsity of connectivity was maintained (details in “Sparsity and rewiring” section).

### 4.3 Time

SNN trials, whether they do or do not involve training, all take place over time. We use a discrete time step *δt* = 1*ms* for all our SNN work. Input (*x*) is given to the SNN ms-by-ms, and output (*y*) is read ms-by-ms. Dynamics such as all neurons’ membrane potentials (*v*) and spikes (*z*) change over ms as well. The current ms time point is denoted as *t*, the next time point as *t* + 1, and so on.

### 4.4 ALIF model neuron

The spiking neuron model used for all units in the recurrent SNN is the adaptive leaky-integrate-and-fire (ALIF) model. ALIF units contain two hidden state variables – one for the membrane potential (aka voltage) *v* and one for the variable *a* which governs the adaptive spike threshold *A*. Together they determine whether a unit *i* spikes (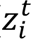 = 1) or does not spike (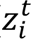 = 0) at time point *t*.

Each unit’s membrane potential evolves over time according to the equation:

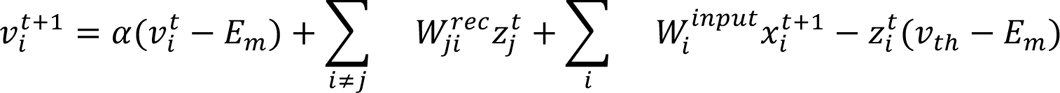

where the resting membrane potential *E_m_*, = −70.6*mV*, the baseline threshold *v*_)._ = −50.4*mV*, the decay factor *a* = *e*^−*δt*/*τm*^, and *τ*_*m*_ = 20*ms* is the membrane time constant.

All synaptic connections and inputs in our model are current-based. At time point *t* + 1, SNN unit *i* receives recurrent spiking input from its presynaptic units *j* which just spiked (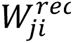, weighted according to 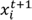), and a subset also receive stimulus input (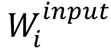, weighted according to 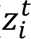). This is summed across all presynaptic units and all stimulus input sources, so that the input current into unit *i* at time *t* + 1 is given as:

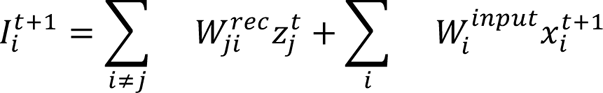

Finally, the term 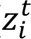(*v_th_-E_m_*) reduces a unit’s membrane potential by a constant value after neuron i spikes (Bellec et al. 2020). 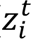 is further fixed to be 0 for a refractory period of 4ms following unit i spiking.

By the above equations, all units’ membrane potentials evolve over time. When a unit’s membrane potential exceeds the spike threshold, that unit will emit a spike. For ALIF neurons, the spike threshold is adaptive, meaning it also evolves over time.

The adaptive threshold increases following a spike and decays exponentially to the baseline threshold *v*_th_. This can be described by the equations:

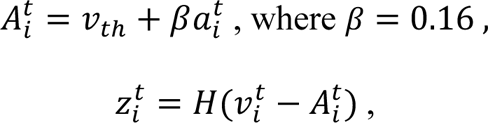

where *H* is the Heaviside step function,

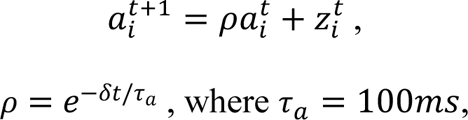

which can be altered to fit timescales relevant to the task of interest (Bellec et al. 2020).

Essentially, if a unit’s membrane potential at time t exceeds its adaptive threshold at time t, the unit will emit a spike.

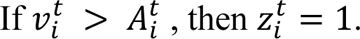

At the very beginning when *t* = 0, voltages are initialized with a random normal distribution (μ = −65 mV, σ = 5 mV).

### 4.5 Structure

SNNs are initialized with neocortical structural properties, some of which continue to be enforced during training. Initial excitatory weights follow a long-tailed, log-normal distribution (Song et al. 2005), where μ = −0.64 nA, σ = .51 nA, corresponding to a mean of 0.6005 nA and a variance of 0.1071 nA (Bojanek & Zhu et al. 2020). Inhibitory weights follow the same distribution but with values 10x stronger than excitatory. All units are sparsely and recurrently connected; the precise probabilities of connection within and between e and i populations are taken from neocortical experiments (Billeh et al. 2020). Weight values and connection probabilities of e and i populations are permitted to change during training. However, overall sparsity is maintained (details in “Sparsity and rewiring” section).

### 4.6 Dynamics

In addition to structural constraints, models are also constrained to exhibit spiking dynamics that match neocortical dynamics during training. For example, spiking in neocortex is sparse, asynchronous, and near-critical (Brunel 2000; Renart et al. 2010; Zerlaut et al. 2019), and each of these features can be maintained in the SNNs’ activity throughout training. Further details are provided in the “Training” section.

### 4.7 Output

Output from the main SNN will vary depending on the task. A subset of main SNN units sends weighted connections to the output. Thus, 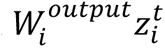 is the output activation from SNN unit *i* at time *t*. Output connectivity is initialized using the same statistics as the recurrent SNN’s connectivity, and sparsity is maintained during training in the same manner.

### 4.8 Training

SNNs are trained using backpropagation-through-time (BPTT) with modifications for spiking. The goal of training is to reduce the difference between the SNNs’ output and the desired output, while still maintaining naturalistic dynamics and structure. This is achieved through iteratively modifying the weights of the recurrent SNN. Input and output weights may also be permitted to change.

Since tasks are temporal, the task error (aka loss) at each timepoint is the mean squared error between the desired output (aka target) at time t and the SNNs’ actual output (aka prediction) at time t.

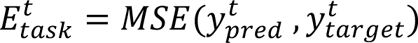

To maintain naturalistic dynamics, additional components are added to the total loss. For example, a target spike rate can be specified. The MSE deviation of the SNN’s actual spike rate from that target rate is added to the loss (Zhu et al. 2020).

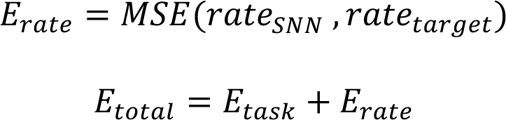

For all our experiments, we train networks in three ways: to minimize task loss, to minimize rate loss, and to minimize both task and rate loss. We report results separately for all three cases for each version of our SNNs.

### 4.9 Gradient descent with Adam optimizer

The goal of training is to reduce the total loss. To do so, we iteratively modify the weights in the network using a version of stochastic gradient descent.

The amount by which each weight should change in each iteration is determined by its gradient, which can be written as:

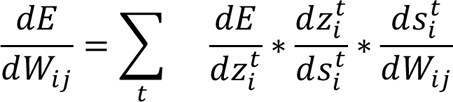

The only new variable here is 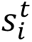, which is the hidden state of neuron *i* at time *t*, and includes membrane potential and adaptation. 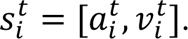.

Since the task and training take place over time, the total loss is summed over all time points *t*. Also, since the network is recurrently connected, the output, the hidden state of all units, and the spiking activity of all units at time *t* is dependent on all other units’ states and activities at time *t* − 1, and at time *t* − 2, and so on to *t* = 0. Those previous states and activities in turn depend on the weights. Therefore, terms are recursively expanded over past time steps (details in Bellec et al. 2020).

We use the Adam optimizer to determine precisely how weights should change based on the gradients. Typically in stochastic gradient descent, each gradient is multiplied by a static learning rate to yield the value by which each weight should change. The Adam optimizer instead adapts the learning rate for every variable over the course of training (details in Kingma & Ba 2014).

### 4.10 Spike pseudo-derivative

The above is the standard form of BPTT for recurrent neural networks. However, because the units in our network spike, and spikes are not differentiable (recall that 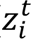 is determined by the Heaviside step function), we use a pseudo-derivative to replace the term 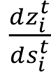 (Huh & Sejnowski 2018, Bellec et al. 2020). Outside of the refractory period, the pseudo-derivative is defined as:

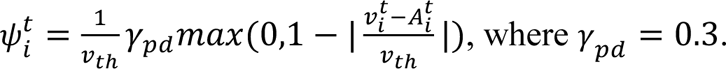

During the refractory period, 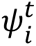.

### 4.11 Sparsity and rewiring

To maintain the overall level of connectivity in the SNN, all weights in all layers that are initialized as 0’s (*W_ij_* = 0 indicates that units *i* and *j* are “disconnected”) will maintain their 0 values during training. All other weights can be updated via gradient descent. An exception occurs when an existing edge weight flips sign (e.g. an excitatory edge weight becomes negative) or becomes 0. In that case, that edge is set to 0 (e.g. is “pruned” away) and a new edge is randomly drawn from the pool of 0-valued edges. The new edge’s weight value is drawn from the initial weight distribution. This process is akin to synaptic rewiring in neocortex (Bellec et al. 2018).

### 4.12 Task

Models were trained on a visual motion entropy change detection task. The model is presented with videos of 460 white drifting dots on a black background, with dots moving in one of 4 global directions (0°, 90°, 180°, 270°), and at two motion entropy levels (low: 100% or high: 15%, which is the percentage of dots that are moving together in the same direction). All dots move with speeds of 10 pixels per ms. Dots are randomly placed on the screen at the start of each video trial and are initialized with random remaining durations to their lifetimes (total lifetime of 1000 ms). Each time a dot reaches its max lifetime, it is removed, and a new dot is randomly drawn at a new location.

Each video trial has a total duration of 4080 ms. Half of all trials have a change in motion entropy which occurs at a random time between 500 and 3500 ms within the trial. The task is for the model to report the motion entropy level at all timepoints of the trial.

### 4.13 CNN front-end

To make the videos interpretable to the main SNN, we created and trained a CNN model of retina and thalamus as a preprocessor that turns videos into spike sequences (Maheswaranathan et al. 2023). This CNN model was trained to report the velocity (x,y) of global motion in a combined dataset of drifting dot videos and black-and-white natural motion videos.

**Figure S1:**
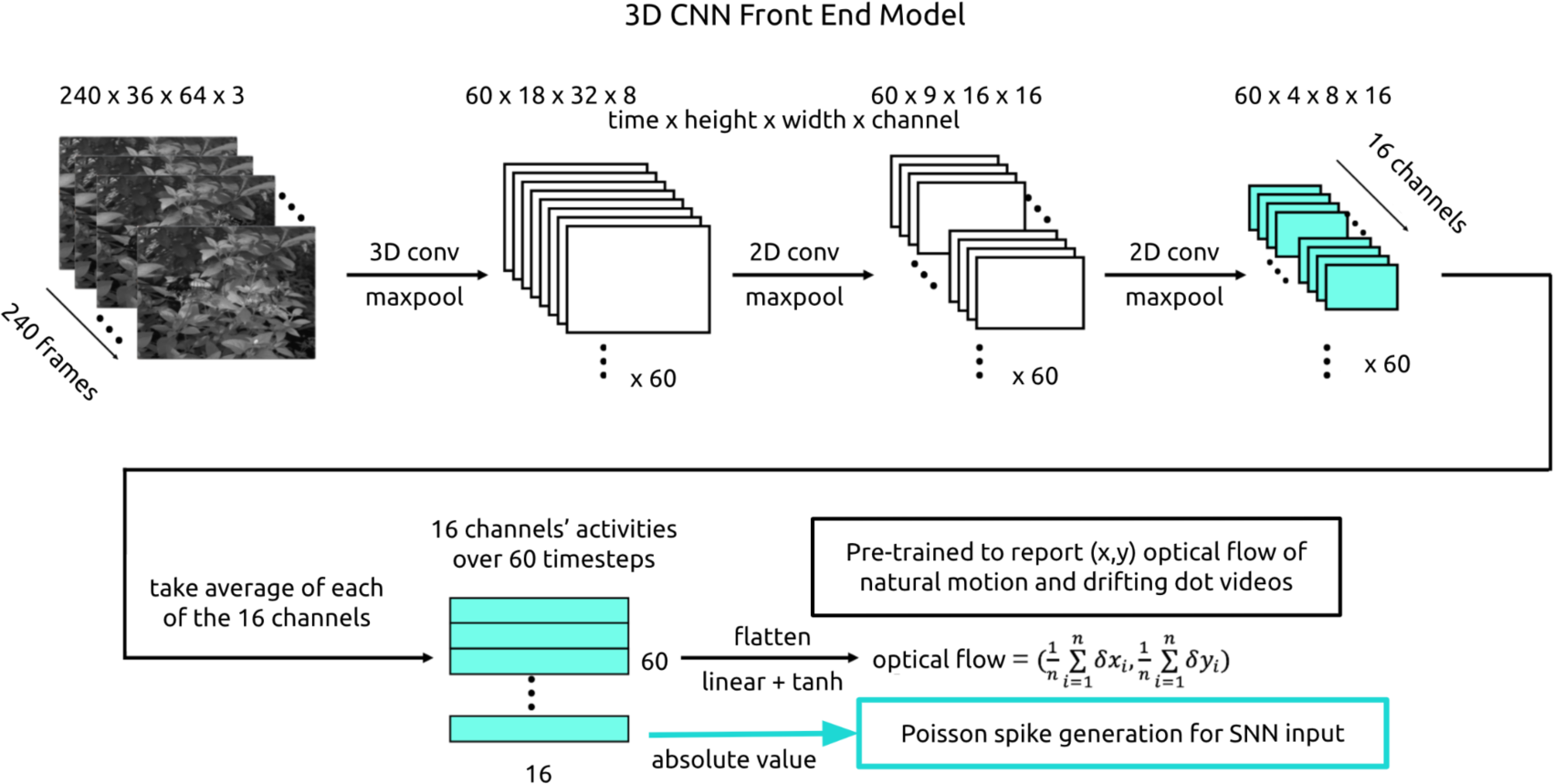
Model front-end architecture. To make videos interpretable to the main SNN, we built a 3D CNN front-end. The absolute value activations of the CNN’s last layer were interpreted as firing rates from which Poisson spikes were generated to become main SNN input.

The CNN model is composed of three successive blocks of Conv3D, MaxPooling3D, and dropout layers, after which the output is flattened into a prediction of x velocity and a prediction of y velocity. We take the average output activations of the 16 units in the last layer to indicate Poisson firing rates. Poisson spikes are newly generated for each trial according to these numbers, and then given as input to a subset of main SNN units.

### 4.14 Training sequence

A single trial involves an input sequence and a target output sequence. The input sequence is composed of Poisson spiking activity from 16 input units. The target sequence is 0’s and/or 1’s - the label attached to each motion entropy level is swapped in different experiments.

SNNs are trained on a large set of these trials. There are a total of 600 unique trials, which are repeated and shuffled to create the desired total training set size. Experiments were run for a duration of 30 trials per batch x 10,000 total batch updates, which yields 300,000 total trials.

To reduce overfitting, in which the SNN becomes overly good at particular trials but not others, we accumulate *E_total_* and 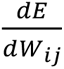 over a batch of 30 trials before updating wesights. The process then repeats for the next batch of 30 trials. Due to trial shuffling and repetition, each batch contains a unique set of trials. In this way, weight changes will improve the average performance over many different trials.

### 4.15 Software and hardware

All SNN training was completed using Python 3.8 or higher run on Nvidia GPUs with CUDA version 11.2 or higher.

### 4.16 Statistical analysis

To compare all distributions of SNN measures, such as naive and trained firing rates to the two motion entropy levels, naive and trained weight distributions of input, recurrent, and output layers, etc., we performed Kolmogorov-Smirnov two sample testing and reported both the distance between means and the p-value.

### 4.17 Weight changes during rate-only and task-only training

A similar pattern of weight changes occurred during training on only rate as during rate-and-task training. Recurrent weights changed the most (e→e: 0.009 ± 0.026 naive to 0.131 ± 0.358 trained; e→i: 0.012 ± 0.028 naive to 0.153 ± 0.413 trained; i→e: −0.150 ± 0.306 naive to 0.531 ± 1.655 trained; i→i: −0.171 ± 0.325 naive to −0.353 ± 1.457 trained; p ≈ 0.0 for all naive and trained recurrent weight distributions). Inputs to excitatory units changed more than inputs to inhibitory units (in→e: 0.225 ± 0.502 naive to 0.313 ± 0.761 trained, p = 1.827 · 10^-40^; in→i: 0.184 ± 0.459 naive to 0.199 ± 0.555 trained, p = 0.030). Output layer weights did not change (e→out: 0.009 ± 0.025 naive to 0.009 ± 0.025 trained, p = 1.0; i→out: −0.150 ± 0.307 naive to −0.150 ± 0.307 trained, p = 1.0).

After training on only task, recurrent weights also changed the most (e→e: 0.010 ± 0.025 naive to 0.126 ± 0.366 trained, p ≈ 0.0; e→i 0.012 ± 0.028 naive to 0.074 ± 0.335 trained, p =2.200 · 10^-298^; i→e −0.151 ± 0.309 naive to −0.656 ± 1.915 trained, p ≈ 0.0; i→i −0.170 ± 0.322 naive to −0.695 ± 1.923 trained, p = 1.319 · 10^-118^), while input layer weights to inhibitory units (in→e: 0.313 ± 0.563 naive to 0.398 ± 0.744 trained, p = 3.137 · 10^-17^; in→i: 0.272 ± 0.533 naive to 0.255 ± 0.576 trained, p = 0.077) and output layer weights (e→out: 0.010 ± 0.027 naive to 0.007 ± 0.022 trained, p = 0.202; i→out: −0.136 ± 0.286 naive to −0.129 ± 0.272 trained, p = 1.0) did not change.

## Acknowledgments

We thank Wolfgang Maass and Guillaume Bellec for providing us with invaluable initial guidance, unique insights, and code examples for us to begin training SNNs. We thank Elizabeth de Laittre, Harold Rockwell, Gabriella Wheeler Fox, and Tarek Jabri for their comments throughout these investigations which greatly improved our analyses. We thank Tarek Jabri, Josh Cruz, and Benton Girdler for their direct contributions to the codebase. We thank Stephanie Palmer, David Freedman, and John Maunsell for their guidance in the development and execution of this work.

## References

Alitto, H. J., & Dan, Y. (2010). Function of inhibition in visual cortical processing. Current opinion in neurobiology, 20(3), 340–346. 10.1016/j.conb.2010.02.012.

Bellec G, Kappel D, Maass W, Legenstein R. Deep rewiring: Training very sparse deep networks. arXiv:1711.05136v5 [cs.NE] [Preprint]. 2018. https://arxiv.org/abs/1711.05136v5.

Bellec, G., Scherr, F., Subramoney, A., Hajek, E., Salaj, D., Legenstein, R., & Maass, W. (2020). A solution to the learning dilemma for recurrent networks of spiking neurons. Nature communications, 11(1), 3625. https://www.nature.com/articles/s41467-020-17236-y.

Billeh, Y. N., Cai, B., Gratiy, S. L., Dai, K., Iyer, R., Gouwens, N. W., … & Arkhipov, A. (2020). Systematic integration of structural and functional data into multi-scale models of mouse primary visual cortex. Neuron, 106(3), 388–403. 10.1016/j.neuron.2020.01.040.

Bojanek, K., Zhu, Y.**, MacLean, J. N. (2020). Cyclic transitions between higher order motifs underlie sustained asynchronous spiking in sparse recurrent networks. PLOS Comput Biol, 16(9), e1007409. 10.1371/journal.pcbi.1007409. *co-first-authors.

Bos, H., Oswald, A. M., & Doiron, B. (2020). Untangling stability and gain modulation in cortical circuits with multiple interneuron classes. bioRxiv, 2020-06. https://www.biorxiv.org/content/10.1101/2020.06.15.148114v2.abstract.

Brette, R. (2015). Philosophy of the spike: rate-based vs. spike-based theories of the brain. Frontiers in systems neuroscience, 151. 10.3389/fnsys.2015.00151.

Brette, R., & Gerstner, W. (2005). Adaptive exponential integrate-and-fire model as an effective description of neuronal activity. Journal of neurophysiology, 94(5), 3637–3642. https://journals.physiology.org/doi/full/10.1152/jn.00686.2005.

Brunel, N. (2000). Dynamics of sparsely connected networks of excitatory and inhibitory spiking neurons. Journal of computational neuroscience, 8, 183–208. https://link.springer.com/article/10.1023/A:1008925309027.

Brunel, N. (2016). Is cortical connectivity optimized for storing information?. Nature neuroscience, 19(5), 749–755. https://www.nature.com/articles/nn.4286.

Buonomano, D. V., & Maass, W. (2009). State-dependent computations: spatiotemporal processing in cortical networks. Nature Reviews Neuroscience, 10(2), 113–125. https://elifesciences.org/articles/73276.

Calaim, N., Dehmelt, F. A., Gonçalves, P. J., & Machens, C. K. (2022). The geometry of robustness in spiking neural networks. Elife, 11, e73276. 10.7554/eLife.73276.

Carandini, M., & Ringach, D. L. (1997). Predictions of a recurrent model of orientation selectivity. Vision research, 37(21), 3061–3071. https://www.sciencedirect.com/science/article/pii/S0042698997001004.

Cohen, U., Chung, S., Lee, D. D., & Sompolinsky, H. (2020). Separability and geometry of object manifolds in deep neural networks. Nature communications, 11(1), 746. https://www.nature.com/articles/s41467-020-14578-5.

Cone, J. J., Scantlen, M. D., Histed, M. H., & Maunsell, J. H. (2019). Different inhibitory interneuron cell classes make distinct contributions to visual contrast perception. Eneuro, 6(1). https://www.ncbi.nlm.nih.gov/pmc/articles/PMC6414440/.

Day-Cooney, J., Cone, J. J., & Maunsell, J. H. (2022). Perceptual weighting of V1 spikes revealed by optogenetic white noise stimulation. Journal of Neuroscience, 42(15), 3122–3132. https://www.jneurosci.org/content/42/15/3122.abstract.

DeAngelis, G. C., Robson, J. G., Ohzawa, I., & Freeman, R. D. (1992). Organization of suppression in receptive fields of neurons in cat visual cortex. Journal of Neurophysiology, 68(1), 144–163. 10.1152/jn.1992.68.1.144.

deCharms, R. C., & Merzenich, M. M. (1996). Primary cortical representation of sounds by the coordination of action-potential timing. Nature, 381(6583), 610–613. https://www.nature.com/articles/381610a0.

Douglas, R. M., Neve, A., Quittenbaum, J. P., Alam, N. M., & Prusky, G. T. (2006). Perception of visual motion coherence by rats and mice. Vision research, 46(18), 2842–2847. 10.1016/j.visres.2006.02.025.

Eysel, U. T., Crook, J. M., & Machemer, H. F. (1990). GABA-induced remote inactivation reveals cross-orientation inhibition in the cat striate cortex. Experimental Brain Research, 80, 626–630. https://link.springer.com/article/10.1007/BF00228003.

Ferster, D., & Miller, K. D. (2000). Neural mechanisms of orientation selectivity in the visual cortex. Annual review of neuroscience, 23(1), 441–471. 10.1146/annurev.neuro.23.1.441.

Huh, D., & Sejnowski, T. J. (2018). Gradient descent for spiking neural networks. Advances in neural information processing systems, 31. http://papers.nips.cc/paper/7417-gradient-descent-for-spiking-neural-networks.

Jabri, T., & MacLean, J. N. (2022). Large-scale algorithmic search identifies stiff and sloppy dimensions in synaptic architectures consistent with murine neocortical wiring. Neural Computation, 34(12), 2347–2373. 10.1162/neco_a_01544.

Kandel, E. R. (1957). Dale’s principle and the functional specificity of neurons. Psychopharmacology; A Review of Progress, 1967, 385–398.

Katzner, S., Busse, L., & Carandini, M. (2011). GABAA inhibition controls response gain in visual cortex. Journal of Neuroscience, 31(16), 5931–5941. https://www.jneurosci.org/content/31/16/5931.short.

Kepecs, A., & Fishell, G. (2014). Interneuron cell types are fit to function. Nature, 505(7483), 318–326. https://www.nature.com/articles/nature12983.

Kingma, D. P., & Ba, J. (2014). Adam: A method for stochastic optimization. *arXiv* preprint arXiv:1412.6980. 10.48550/arXiv.1412.6980.

Kirkels, L. A. M. H., Zhang, W., Havenith, M. N., Tiesinga, P., Glennon, J., Van Wezel, R. J. A., & Duijnhouwer, J. (2018). The opto-locomotor reflex as a tool to measure sensitivity to moving random dot patterns in mice. Scientific reports, 8(1), 1–9. https://www.nature.com/articles/s41598-018-25844-4.

Kohn A, Coen-Cagli R, Kanitscheider I, Pouget A. (2016). Correlations and neuronal population information. Annual review of neuroscience, 39: 237–256. 10.1146/annurev-neuro-070815-013851.

Koulakov, A. A., Hromádka, T., & Zador, A. M. (2009). Correlated connectivity and the distribution of firing rates in the neocortex. Journal of Neuroscience, 29(12), 3685–3694. 10.1523/JNEUROSCI.4500-08.2009.

Lagzi, F., Bustos, M. C., Oswald, A. M., & Doiron, B. (2021). Assembly formation is stabilized by Parvalbumin neurons and accelerated by Somatostatin neurons. bioRxiv, 2021-09. https://www.biorxiv.org/content/10.1101/2021.09.06.459211v1.abstract.

Lankarany M, Prescott SA. (2015). Multiplexed coding through synchronous and asynchronous spiking. BMC Neuroscience, 16(1): 1–2. 10.1186/1471-2202-16-S1-P198.

Lee, J. H., Delbruck, T., & Pfeiffer, M. (2016). Training deep spiking neural networks using backpropagation. Frontiers in neuroscience, 10, 508. https://www.frontiersin.org/articles/10.3389/fnins.2016.00508/full.

Litwin-Kumar, A., & Doiron, B. (2012). Slow dynamics and high variability in balanced cortical networks with clustered connections. Nature neuroscience, 15(11), 1498–1505. https://www.nature.com/articles/nn.3220.

Maheswaranathan, N., McIntosh, L. T., Tanaka, H., Grant, S., Kastner, D. B., Melander, J. B., … & Baccus, S. A. (2023). Interpreting the retinal neural code for natural scenes: From computations to neurons. Neuron.

Marques, T., Summers, M. T., Fioreze, G., Fridman, M., Dias, R. F., Feller, M. B., & Petreanu, L. (2018). A role for mouse primary visual cortex in motion perception. Current Biology, 28(11), 1703–1713. 10.1016/j.cub.2018.04.012.

Mejias, J. F., & Longtin, A. (2012). Optimal heterogeneity for coding in spiking neural networks. Physical Review Letters, 108(22), 228102. https://journals.aps.org/prl/abstract/10.1103/PhysRevLett.108.228102.

Morrison, D. J., Rashid, A. J., Yiu, A. P., Yan, C., Frankland, P. W., & Josselyn, S. A. (2016). Parvalbumin interneurons constrain the size of the lateral amygdala engram. Neurobiology of learning and memory, 135, 91–99. 10.1016/j.nlm.2016.07.007.

Morrone, M. C., Burr, D. C., & Maffei, L. (1982). Functional implications of cross-orientation inhibition of cortical visual cells. I. Neurophysiological evidence. Proceedings of the Royal Society of London. Series B. Biological Sciences, 216(1204), 335–354. https://royalsocietypublishing.org/doi/abs/10.1098/rspb.1982.0078.

Renart, A., De La Rocha, J., Bartho, P., Hollender, L., Parga, N., Reyes, A., & Harris, K. D. (2010). The asynchronous state in cortical circuits. Science, 327(5965), 587–590. https://www.science.org/doi/full/10.1126/science.1179850.

Resulaj, A., Ruediger, S., Olsen, S. R., & Scanziani, M. (2018). First spikes in visual cortex enable perceptual discrimination. Elife, 7, e34044. https://elifesciences.org/articles/34044.

Roxin, A., Brunel, N., Hansel, D., Mongillo, G., & van Vreeswijk, C. (2011). On the distribution of firing rates in networks of cortical neurons. Journal of Neuroscience, 31(45), 16217–16226. https://www.jneurosci.org/content/31/45/16217.short.

Sharmin, S., Rathi, N., Panda, P., & Roy, K. (2020). Inherent adversarial robustness of deep spiking neural networks: Effects of discrete input encoding and non-linear activations. In Computer Vision–ECCV 2020: 16th European Conference, Glasgow, UK, August 23–28, 2020, Proceedings, Part XXIX 16 (pp. 399–414). Springer International Publishing. https://link.springer.com/chapter/10.1007/978-3-030-58526-6_24.

Shew, W. L., Yang, H., Petermann, T., Roy, R., & Plenz, D. (2009). Neuronal avalanches imply maximum dynamic range in cortical networks at criticality. Journal of neuroscience, 29(49), 15595–15600. 10.1523/JNEUROSCI.3864-09.2009.

Song, S., Sjöström, P. J., Reigl, M., Nelson, S., & Chklovskii, D. B. (2005). Highly nonrandom features of synaptic connectivity in local cortical circuits. PLoS biology, 3(3), e68. 10.1371/journal.pbio.0030068.

Verzi, S. J., Rothganger, F., Parekh, O. D., Quach, T., Miner, N. E,, Vineyard, C. M., James, C. D., Aimone, J. B. (2018). Computing with spikes: The advantage of fine-grained timing. Neural Computation, 30, 2660–2690. 10.1162/neco_a_01113.

Zenke, F., & Vogels, T. P. (2021). The remarkable robustness of surrogate gradient learning for instilling complex function in spiking neural networks. Neural computation, 33(4), 899–925. 10.1162/neco_a_01367.

Zerlaut, Y., Zucca, S., Panzeri, S., & Fellin, T. (2019). The spectrum of asynchronous dynamics in spiking networks as a model for the diversity of non-rhythmic waking states in the neocortex. Cell reports, 27(4), 1119–1132. 10.1016/j.celrep.2019.03.102.

Zhu, Y., Scherr, F., Maass, W., MacLean, J. (2020, November 9-12). Addition of neocortical features permits successful training of spiking neuronal network models [Conference presentation]. From Neuroscience to Artificially Intelligent Systems, Cold Spring Harbor Laboratory, NY, United States. https://meetings.cshl.edu/meetings.aspx?meet=naisys&year=20.

